# SnakeChunks: modular blocks to build Snakemake workflows for reproducible NGS analyses

**DOI:** 10.1101/165191

**Authors:** Claire Rioualen, Lucie Charbonnier-Khamvongsa, Jacques van Helden

## Abstract

**Summary:** Next-Generation Sequencing (NGS) is becoming a routine approach for most domains of life sciences, yet there is a crucial need to improve the automation of processing for the huge amounts of data generated and to ensure reproducible results. We present SnakeChunks, a collection of Snakemake rules enabling to compose modular and user-configurable workflows, and show its usage with analyses of transcriptome (RNA-seq) and genome-wide location (ChIP-seq) data.

**Availability:** The code is freely available (github.com/SnakeChunks/SnakeChunks), and documented with tutorials and illustrative demos (snakechunks.readthedocs.io).

**Supplementary information:** Supplementary data are available at *Bioinformatics* online.

## 1 Introduction

Beyond the huge increase in computer resources required by the rapid expansion of Next-Generation Sequencing technologies, the community recognizes the urgent need for software systems ensuring the reproducibility of the analytic workflows. Indeed, although any publication is conditioned to the deposit of raw sequences in short read archives, there is still no incentive to provide a formal description that would enable a third-party to reproduce the full analytic workflow or even trace the steps and parameters that led from the raw sequences to the published results.

Snakemake (Köster and Rahmann, 2012) is a flexible workflow engine developed for large-scale biological data analysis, which is being adopted by a growing community of users. The common practice is to implement a separate script for each analysis, specifying the commands for each step of the workflow (rules) and the desired output files (targets).

Building up on the Snakemake concepts, we have developed SnakeChunks, a collection of modular rules, which can be used either in predefined workflows, or combined to compose custom pipelines. They are portable to different operating systems, and allow the automation of the operations, and traceability of the tools and parameters used. We demonstrate their usage with workflows to analyse transcriptome (RNA-seq) and genome-wide location (ChIP-seq) data.

## 2 SnakeChunks

### 2.1 Rules and workflows

Snakemake is a Python library inspired by concepts of the *make* language:

- rules are recipes to perform an operation (using python, R or shell);
- dependencies between rules are inferred by inputs and outputs names;
- expected result files are defined by a list of targets.

The SnakeChunks library currently contains 70 rules, organised in separate files to enable their modular assembly into workflows. This flexibility of composition is illustrated by 5 demo workflows. A given rule can thus be recycled in several workflows, avoiding code duplication, and can readily be included in new custom pipelines.

We have interfaced a variety of alternative tools for crucial steps such as download from short read databases, read mapping, peak-calling for ChIPseq, differential expression analysis for RNA-seq. For each one of these steps, one or several alternative tools can be selected in the configuration file. Taking benefit of Snakemake’s capacity of parallelization, this modularity eases tool benchmarking and parameter optimization.

The workflows produce various types of reports and outputs: flowcharts of the analyses, quality reports (MultiQC, FastQC), differential expression analysis report (SARTools), IGV session, RSAT peak-motifs report (see Supplementary figures).

**Fig 1.**
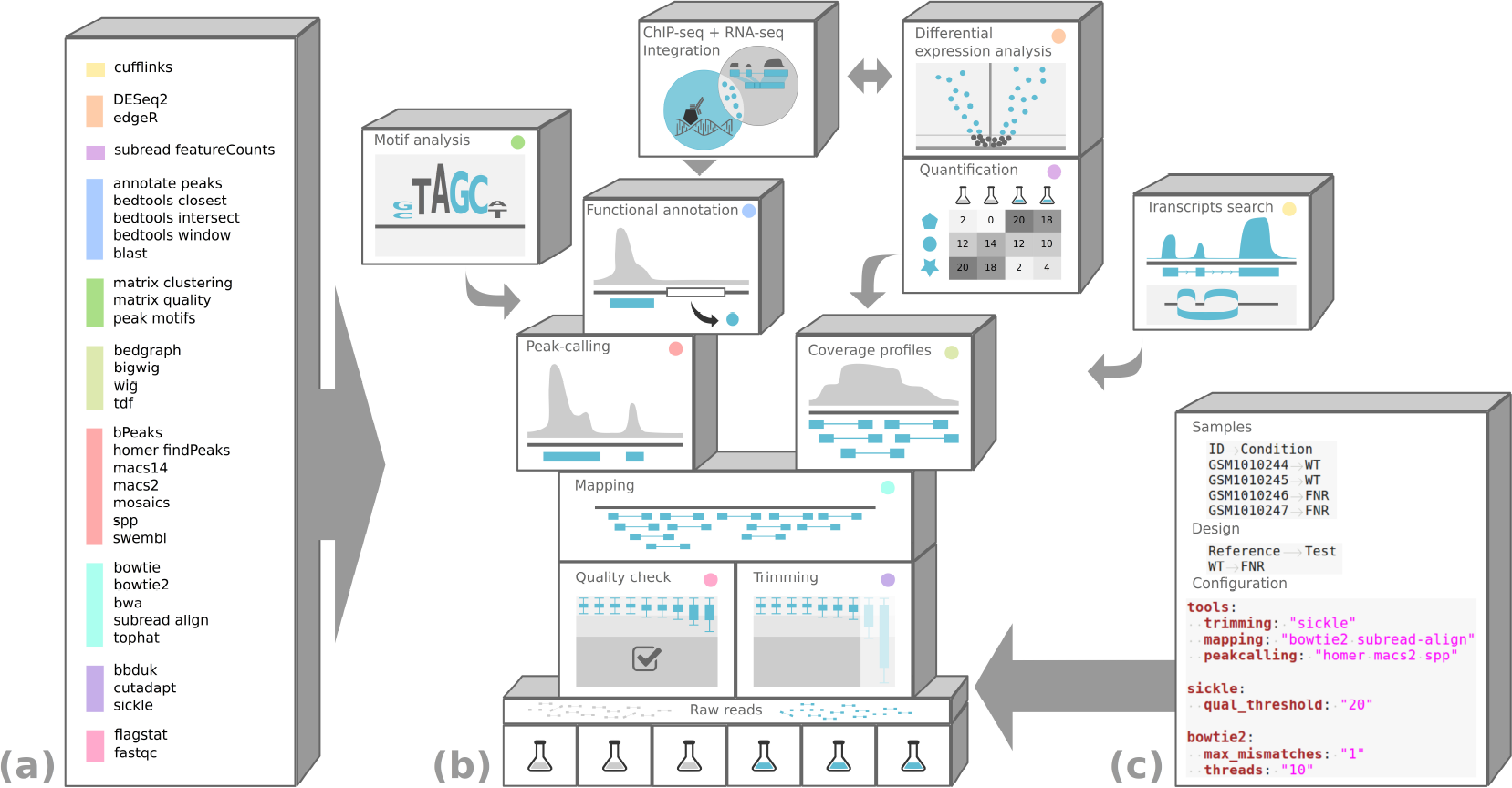
SnakeChunks framework. A variety of rules (a) can be assembled to create custom workflow structures (b), whose metadata and parameters are specified in independent files (c).

### 2.2 Metadata and parameters

Different tools can be chosen for each step, and their parameters need to be fine-tuned to cater workflows to different datasets. Our generic workflows can be adapted to a given study case by editing three files:

- a tab-delimited file containing the list of samples to process, as well as any relevant information about these samples;
- a tab-delimited file describing the design of the analysis (*e.g*. conditions or samples to be compared);
- a YAML-formatted configuration file, specifying the analysis-specific information: directories, reference genome, annotation files, tools, and optionally, customized parameters for each tool.

These metadata and parameter files actually contain all the required information to submit NGS data to short read databases.

### 2.3 Tutorials

Several tutorials are available in the documentation (snakechunks.readthedocs.io), to facilitate the use of the code.

- Introduction to Snakemake key concepts, with practical exercices.
- Execution of SnakeChunks workflows on publicly available data.
- Execution of SnakeChunks workflows on user-provided data, and customization with the configuration file.
- Installation of all the dependencies organized in thematic sections and described tool by tool.
- Creation of virtual machines to keep a working trace of the programs used and their versions.
- Execution of a demo in a Docker container.

## 3 Conclusion and Perspectives

The efficiency and flexibility of the SnakeChunks framework was demonstrated with a variety of use cases of RNA-seq and ChIP-seq analysis in *D. melanogaster*, *A. thaliana* (Castro-Mondragón, Rioualen *et al*., 2016), mouse, yeast, bacteria…In a study of *Glossina palpalis* infection by *Trypanosoma congolense* (Tsagmo *et al*., 2017), we submitted the raw data in GEO with an archive containing the version of the SnakeChunks library used to produce the results along with the metadata and configuration files, which enables anyone to reproduce the full processing from the raw reads to the published results.

All of this material is versioned and available in a public GitHub repository (github.com/SnakeChunks/SnakeChunks). It has been successfully ported and tested on unix servers, Linux and Mac OS X personal computers, and in virtual environments. A Docker image is available (hub.docker.com/r/snakechunks/snakechunks).

The SnakeChunks library is progressively enriched with new rules to incorporate additional tools. It also constitutes a suitable framework to develop new workflows dedicated to other type of NGS analyses, such as variant calling or metagenomics, fields of growing interest for their potential applications in health and ecology.

One of the main impediments for the generalization of Snakemake workflows probably lies in the fact that potential users with a background in life sciences might be reluctant to use command line-based tools. In order to overcome this major obtacle, a GUI was developed (Desvillechabrol *et al*., 2017), that allows to use predefined pipelines or custom pipelines, and edit the configuration directly via the interface.

## Acknowledgements

This work was supported by France Génomique [ANR-10-INBS-09-10]; and National Institute of Health [grant GM0110597] & FOINS-CONACYT - Fronteras de la Ciencia 2015 - ID 15. We thank Thomas Cokelaer for his collaboration and advice.

